# Acute Sprint Exercise Transcriptome in Human Adipose Tissue: Associations with Growth Hormone

**DOI:** 10.64898/2026.06.01.728459

**Authors:** Mona Esbjörnsson, Håkan C Rundqvist, Barbara Norman, Ted Österlund, Jens Bülow, Eva Jansson

## Abstract

It was hypothesised that sprint exercise induces changes in adipose tissue (AT) gene expression related to lipolysis, and that growth hormone (GH) acts as a stimulus. Twelve healthy males and females perform 3×30-s all-out cycle sprints (SIT) and six unloaded cycling (CON). AT biopsies were performed pre and 2 hours post-exercise. Serum GH response was greater in SIT than CON, but no enrichment of differentially expressed genes (DEGs) related to lipolysis was found in AT. However, the GH-receptor was one of few DEGs in AT with interaction between SIT and CON and exercise-induced changes in expression of the pre-selected GH-related targets, *CISH, PTEN* and *G0S2* were associated with exercise-induced increase in GH. This supports a GH-mediated effect of SIT on lipolysis. In a genome-wide analysis, exercise-induced changes in expression of *ARHGAP24, TAF4B, ARID5B, MIR604* and *MIR938*, were associated with increase in GH. The function of these genes is not well understood, but some relationship to AT has earlier been demonstrated Sex influenced GH-response to exercise and expression of the GH-responsive gene *CISH*. Among the most downregulated genes were the core clock genes *PER1, NR1D1* and *CIART*, which decreased in both SIT and CON. This underscores the necessity to control for diurnal variations and sex in future exercise studies on regulation of AT lipolysis.

## INTRODUCTION

Sprint interval training (SIT) has been shown to reduce body fat mass or waist circumference (1, 2), despite the low energy expenditure during a SIT session (3, 4). This may be explained by an increased rate of post-exercise lipolysis and the thermogenic triglyceride-free fatty acid (TG-FFA) cycle, resulting in increased post-exercise energy expenditure (4-6). Acute SIT is known to increase plasma catecholamines, growth hormone (GH), and IL-6 (7, 8), all potential candidates for increased lipolysis and TG-FFA cycle related thermogenesis in adipose tissue (AT) during post-exercise recovery (9-14). Measurements of plasma glycerol from blood draining subcutaneous abdominal AT have confirmed that SIT stimulates lipolysis in adipose tissue (8).

Recently, we tested the hypothesis that post-exercise expression of genes related to fat and energy metabolism would increase in response to SIT. However, this hypothesis was not confirmed by evaluating AT biopsies performed 15 minutes after the last sprint. On the contrary, the expression of such genes decreased (15).Notably, GH has a delayed lipolytic effect, demonstrated both at rest after a bolus injection of GH (16) and post-exercise, with lipolysis starting around one to two hours post-exercise (12). Moreover, GH increases the phosphorylation of the transcription factor STAT5 in AT, in a GH-receptor (GHR) dependent manner (16), and subsequently also the expression of GH-responsive genes (10, 17, 18). Consequently, increases in the expression of genes related to fat and energy metabolism post- \exercise may occur at a later time point than the 15 minutes previously studied (15).

To capture this potential delayed GH-mediated response, AT biopsies were performed 2 hours post-exercise in the present study. We included both men and women in equal proportions, as sex-related differences in GH release and GH-responsive genes have been reported (19-23). For example, circulatory GH shows a faster early response to exercise in women than in men (7, 20, 24-27). Furthermore, a control group of both sexes was included to adjust for other drivers of gene expression in AT besides exercise, such as diurnal (circadian) rhythm (28).

It was hypothesised that acute SIT induces an increase in post-exercise gene expression in AT related to lipolysis and energy expenditure. This occurs at the time of the GH-induced increase in lipolysis, approximately 2 hours post-exercise. A secondary hypothesis was therefore that exercise-induced changes in expression of GH-responsive genes are related to the increase in serum GH during the acute SIT session.

## METHODS

### Subjects

A total of 18 healthy voluntary subjects (9 females and 9 males) participated in the study. The mean ± SD for age, height, body weight, and body mass index are presented in Table 1. Subjects were recruited from voluntary students enrolled in various study programmes at Karolinska Institutet. Inclusion criteria were healthy, non-smoking young adults of both sexes who engaged in regular physical activity, but not at an elite or competitive level. Individuals taking regular medication, except oral contraceptives, and female subjects in the menstrual phase were excluded from the study. An activity index (29) with values ranging from 5.5 to 20.5 was calculated to estimate physical activity during leisure time; the mean and range for our subjects were 16.5 (11–19.5). The percentage of body fat, fat-free body mass, and fat mass was estimated from skinfold measurements of the triceps, biceps, and subscapular regions (30), and are presented in Table 1. Subjects were fully informed about the procedures and potential risks before providing written informed consent. The study was approved by the Ethics Committee at Karolinska Institutet (Dnr 2015/574-31, 2015/1054-32, 2015/1646-32, 2015/2340-32).

**Table 1.**
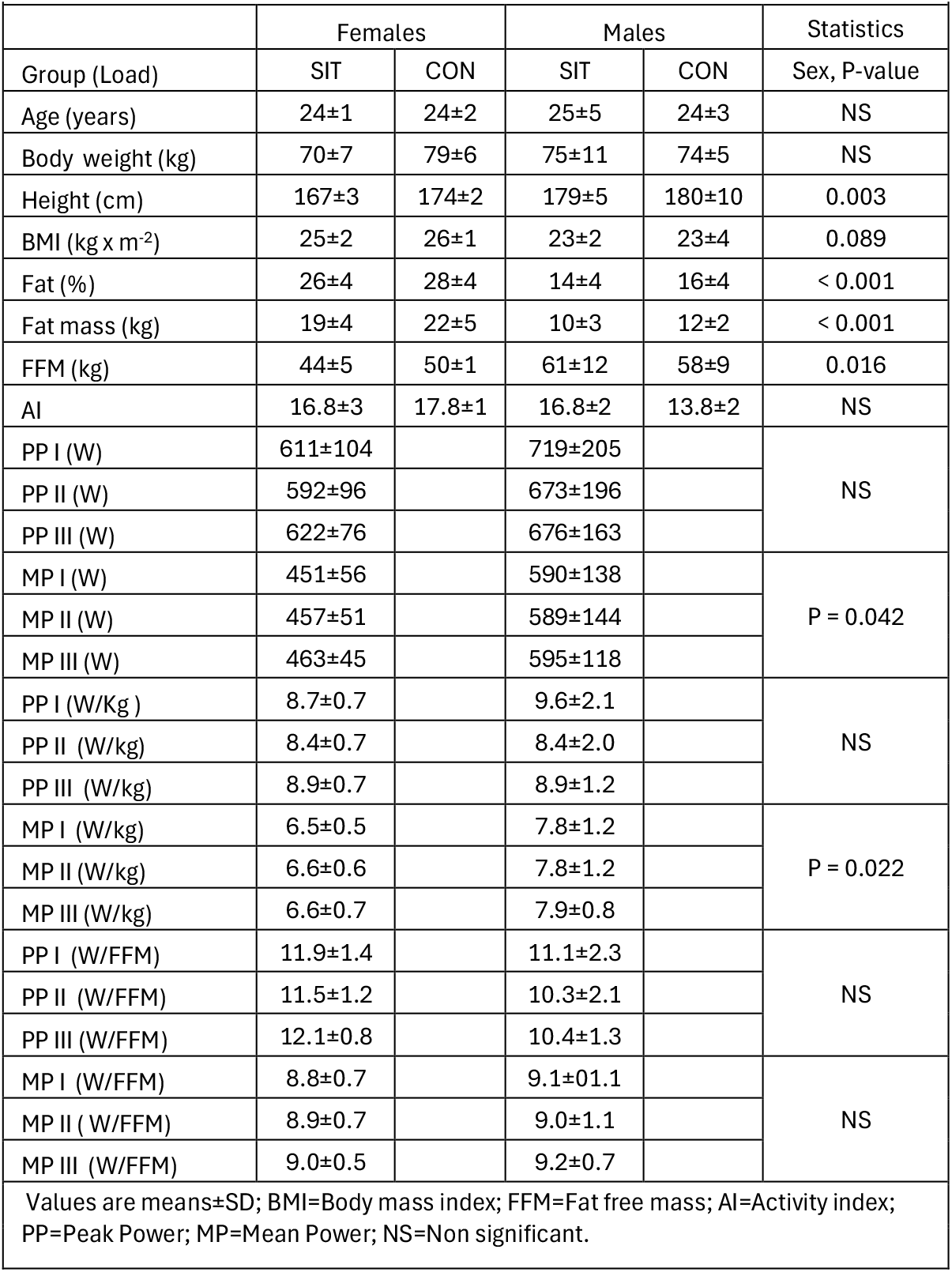
Subject characteristics in SIT (6 females and 6 males; all-out sprints) and CON (3 females and 3 males; unloaded exercise).

### Experimental protocol

After a 60-s warm-up at a cycle ergometer at a load of 25% of the individual braking load at 60 rpm, subjects were randomised to perform repeated bouts of sprint exercise (SIT group; 6 females and 6 males) or unloaded exercise (CON group; 3 females and 3 males). Subjects were asked to refrain from strenuous exercise for 24 hours before the experiment. They reported to the laboratory between 08:00 am and 09:30 after an overnight fast, except for the instruction to consume one to two slices of white bread and a glass of water one hour before arrival. An indwelling catheter was inserted into an antecubital vein. Fifteen mL of blood was sampled in the supine position before a 60-s warm-up, 9 minutes after each bout of exercise, and up to 2 hours after the last bout of exercise (Figure 1). Subcutaneous abdominal adipose tissue biopsies were performed under local anaesthesia without adrenaline, using a Hepafix needle (Braun Medical, Melsungen, Germany): before bout 1 and 2 hours after the last bout of exercise, on different sides of the umbilicus.

**Figure 1.**
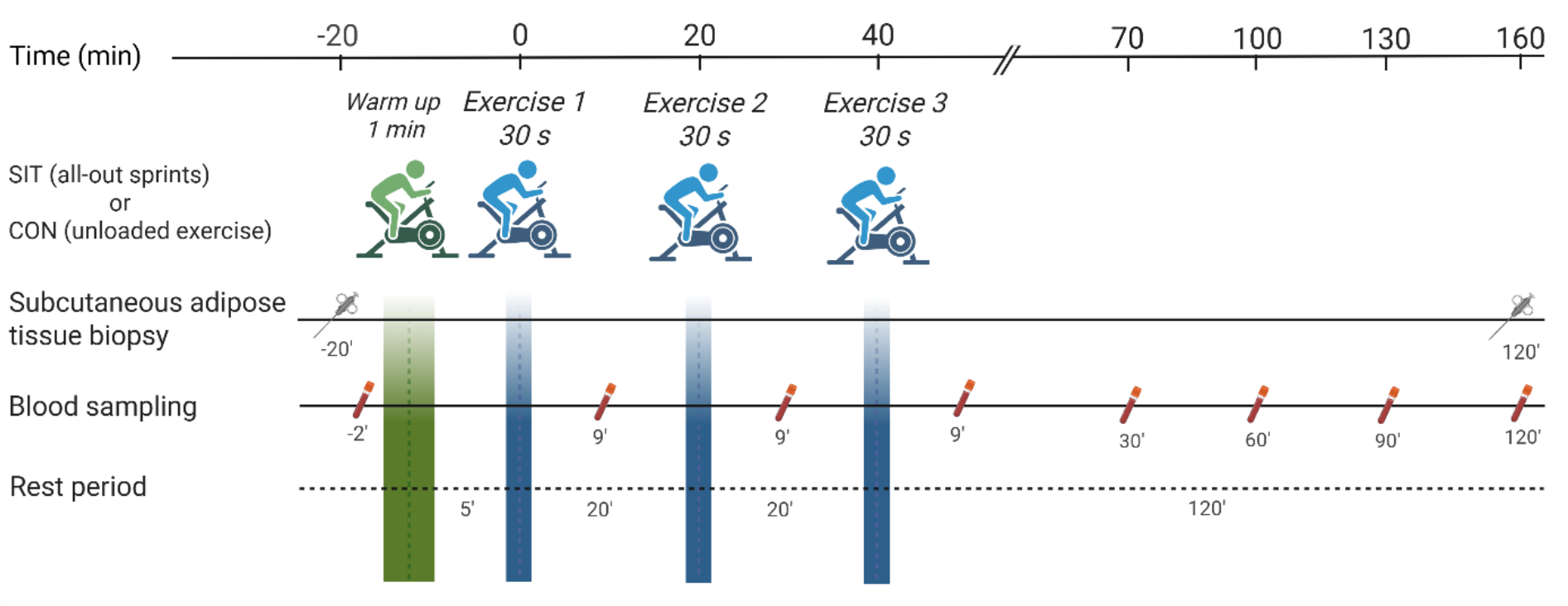
Schematic overview of the experimental protocol. Subjects were randomised after warm up to perform either 3 × 30-s all-out sprints (SIT group; 6 females and 6 males) or unloaded exercise (CON group; 3 females and 3 males). Created in BioRender. Rundqvist, H. (2026) https://BioRender.com/xkmfi67

Subcutaneous abdominal adipose tissue blood flow was measured at rest and up to 2 hours after the last sprint in 3 healthy, physically active subjects, using the same exercise protocol as in the SIT group in the current study (2 females and 1 male, age 20–25 years, BMI 19–28 kg x m^−2^), by the Xenon-133 washout technique (31). The study was approved by the Ethics Committee at Karolinska Institutet (Dnr 2015/574-31, 2015/1054-32, 2015/1646-32).

### Exercise protocol

Subjects in the SIT group performed 3 × 30-s all-out cycle sprints (32) on a mechanically braked cycle ergometer (Monark Ergomedic 894 E, Sweden), with 20 minutes of rest between sprints. They were instructed to pedal as fast as possible at a load of 0.075 kilopond × kg body mass^−1^. Peak power (the highest 5-s power) and mean power (the average power during the 30-s cycle) were calculated for each of the three 30-s cycle sprints. A 20-minutes recovery period was used to ensure near-complete restoration of power output between sprint bouts, as previously applied in our sprint exercise studies (15). Subjects in the CON group performed 3 × 30-s bouts of unloaded exercise on the cycle ergometer with 20 minutes of rest between each bout and were instructed to pedal at 60 rpm.

### Adipose tissue biopsy preparation and analysis

Subcutaneous adipose tissue samples obtained before and after exercise were immediately blotted, placed on a plastic filter (Sefar Nitex 06-210/233, Bigman AB, Sweden), rinsed in sterile saline, cut into smaller pieces, and frozen in liquid nitrogen. Samples were stored at -80 °C until processed. Frozen tissue samples (30–100 mg) were homogenised with a Polytron homogeniser dispenser (Kinematica AG, Lucerne, Switzerland), and RNA was isolated with a Qiagen RNeasy Lipid Mini Kit (Cat. No. 74804, Thermo Fisher Scientific Inc., Waltham, MA, USA), treated with DNase (Qiagen RNase-Free DNase Set, Cat. No. 79254, Thermo Fisher Scientific Inc., Waltham, MA, USA) and quantified using a Nanodrop spectrometer (ND-1000: NanoDrop®, Thermo Fisher Scientific, Wilmington, DE, USA).Total RNA quality was assessed with an Agilent 2200 Tapestation (Agilent Technologies, Santa Clara, CA, USA). RNA integrity (RIN) values indicated consistent and acceptable RNA quality among the samples, with a median RIN value (range) of 7.7 (5.1–8.4).

Microarray analysis was performed using the Human Gene 2.1 ST Array (Thermo Fisher Scientific Inc., Waltham, MA, USA). 150 ng of total RNA was used to generate amplified sense strand cDNA (sscDNA) targets with the GeneChip® WT Plus Reagent Kit (Thermo Fisher Scientific Inc.). 5.2 µg of fragmented and labelled sscDNA target was hybridised to the Human Gene 2.1 ST Array WT, followed by washing, on a GeneTitan MC instrument (Thermo Fisher Scientific), according to the manufacturer’s protocol. Scanning was performed with the Affymetrix Gene Chip Scanner 3000 7G. Raw intensity data (CEL files) were processed using Affymetrix Transcriptome Analysis Console (TAC) Software, including the Signal Space Transformation – Robust Multi-Chip Analysis (SST-RMA) algorithm.

Differentially expressed genes in response to exercise for SIT and CON were identified by *limma* (33). For the analysis of interaction in response to exercise between SIT and CON robust regression from *limma* was used. QIAGEN’s Ingenuity® Pathway Analysis (IPA®, QIAGEN, Redwood City, CA, USA) was used to identify significantly enriched pathways and upstream regulators from the list of differentially expressed genes. The Human Gene 2.1 ST array was selected as the reference. Regression analysis was performed between post-exercise gene expression and GH peak, adjusted for measurements at rest.

### Blood preparation and analyses

Blood samples were divided into two portions. One portion was transferred into sodium-heparinised tubes and immediately centrifuged at 2,000 g (4 °C) for 10 minutes. One-millilitre aliquots of this plasma were frozen in liquid nitrogen and stored at -80 °C. The other portion was transferred to a serum tube, stored at room temperature for 20 minutes, and then treated in the same way as the heparinised blood. Plasma lactate was analysed using a Radiometer ABL 800 Flex blood gas analyser (Berman & Beving Lab, Triolab, Gothenburg, Sweden), and plasma glucose by the hexokinase method with a Roche/Hitachi cobas c-system (Roche Diagnostics GmbH, Mannheim, Germany). Serum free fatty acids (FFA) were analysed by an enzymatic colorimetric assay (FUJIFILM Wako Chemicals Europe GmbH, NEFA-HR (2), Nordic Biolabs, Täby, Sweden), serum insulin by an electrochemiluminescence immunoassay (Cobas e602/e801, Roche Diagnostics GmbH, Germany), and serum growth hormone (GH) by the IMMULITE 2000 Growth hormone (hGH) chemiluminescent enzyme immunoassay (Siemens Healthcare Diagnostics Products Ltd, United Kingdom).

### Statistics

Most variables were normally distributed, except for serum GH, which was log2-transformed to reduce skewness. Gene expression data were also log2-transformed. Values in the text are given as mean and SD unless otherwise stated. P-values were considered statistically significant at P < 0.05. Exercise-induced changes in blood and gene expression data were analysed using 3-factor ANOVAs: time, loading condition (SIT or CON), and sex. Depending on the outcome of these ANOVAs, 1-factor or 2-factor ANOVAs were performed as sub-analyses (see Results). Student’s t-test for paired observations or groups was used when appropriate. Statistical analysis of the relationship between gene expression data and metabolic or hormonal variables was performed using multiple regression analysis or Pearson’s linear regression analysis.

## RESULTS

### Power development

Peak power (PP) and mean power (MP) for the three bouts in females and males in the SIT group are shown in Table 1. PP and MP did not change significantly across the three sprints. PP and PP per kg body mass did not differ between sexes, but MP and MP per kg body mass were significantly higher in males than in females, regardless of the three sprints.

### Metabolic measurements

Exercise-induced changes in blood data were analysed using 3-factor ANOVAs (time, loading condition, and sex). Interactions between time and load were observed for lactate (P < 0.001), glucose (P < 0.001), and insulin (P = 0.010), but no interactions or main effects of sex were found (Figure 2a and 2b). These blood data were further analysed using 1-factor ANOVAs (time), separated by loading conditions, SIT or CON. For free fatty acids (FFA), an interaction between all three factors was identified (P = 0.012), and data were further analysed using 2-factor ANOVAs (time and sex), separated by loading condition. For GH, two interactions were found in the 3-factor ANOVA: time x load (P = 0.004) and time x sex (P = 0.015). For further GH sub-analyses, see below.

**Figure 2.**
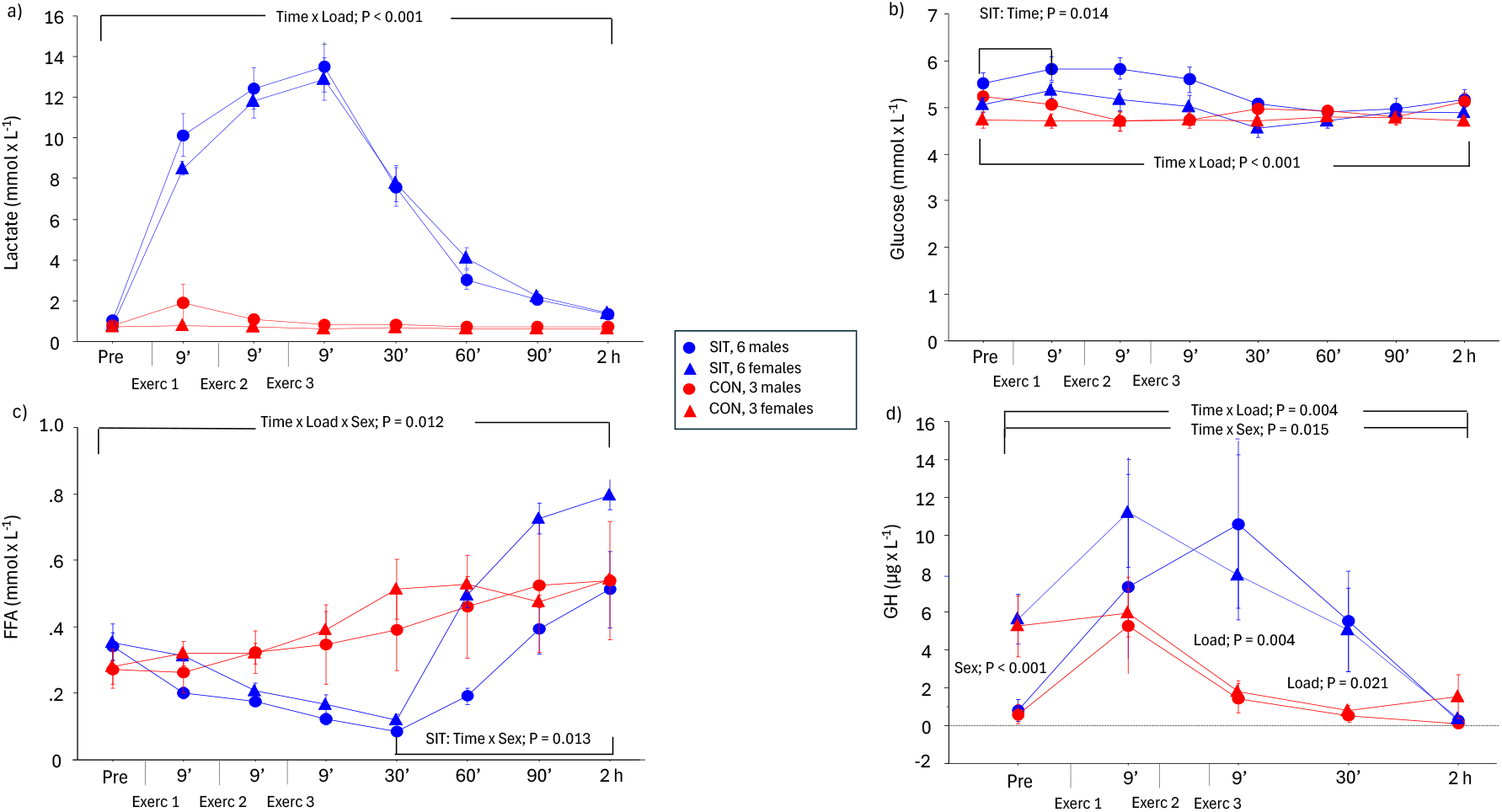
a) Plasma lactate, b) plasma glucose, c) plasma FFA and d) serum GH, pre-exercise, during the exercise session (3 × 30-s with 20 minutes rest in between) and 2 hours after the last bout of exercise in SIT (sprint exercise; 6 males, 6 females) and CON (unloaded exercise; 3 males, 3 females). Data were analysed by 3-factor ANOVAs (time, load and sex) and by 1- or 2-factor ANOVAs for further analysis Serum GH data were log2-transformed. Values are mean ± SE.

Plasma lactate increased in the SIT group during the exercise session and then returned towards baseline (1-factor ANOVA time, P < 0.001). No significant change over time was found in the CON group (Figure 2a).

Plasma glucose increased in the SIT group by 6% between rest and 9 minutes after sprint 1 (P = 0.014), then gradually returned to baseline (1-factor ANOVA time, P < 0.001). No significant change over time was found in CON (1-factor ANOVA time, P = 0.150) (Figure 2b). Plasma FFA decreased in SIT by 60% during exercise (up to 30 minutes post sprint 3), independent of sex (2-factor ANOVA time and sex, P < 0.001 for time). After the exercise session, FFA increased in SIT, with a greater increase in females than males (2-factor ANOVA time and sex, P = 0.013 for time x sex, Figure 2c). In CON, plasma FFA increased over the exercise session, including the post-exercise period, independent of sex (2-factor ANOVA time and sex, P < 0.002 for time).

The total exposure of serum GH during the exercise session, including the 2 hours post-exercise period, differed between SIT and CON, and also between males and females (3-factor ANOVA: time, load, and sex; P = 0.004 for time x load and P = 0.015 for time × sex) (Figure 2d). The time x load interaction for GH was most likely due to total GH exposure being higher in SIT than CON, as GH peak tended to be higher in SIT than CON (P = 0.068,t-test; P = 0.019, Mann-Whitney U test), followed by a faster decline in GH over time in CON than SIT, resulting in approximately 4-fold lower GH levels for CON than SIT at 9 and 30 minutes post third bout of exercise (P = 0.004 and P = 0.021). The time x sex interaction for GH was likely due to GH being 7-fold higher in females than males at rest, independent of loading conditions (2-factor ANOVA: sex and load; P < 0.001 for sex). Females also reached the GH peak earlier than males. In SIT, GH peak was reached after sprint 1 in females and after sprint 3 in males (2-factor ANOVA: time and sex; P < 0.031 for time x sex) (note, GH was not analysed after the second bout of exercise). In CON, the GH peak was reached after the first exercise bout (unloaded exercise) in males, and in females, already at rest. A significant association was found between GH peak and FFA 2 hours after the last sprint (R^2^= 0.48, P = 0.001) (Figure S1).

Serum insulin increased 3-fold during the exercise session (measured only at rest and 2 hours post sprint 3) in SIT (8 ± 2 vs 24 ± 13 mIE x L^−1^; paired t-test, P = 0.001). No significant change was found in CON (7 ± 3 vs 6 ± 2 mIE × L^−1^; paired t-test, P = 0.160).

### Subcutaneous adipose tissue blood flow

Subcutaneous adipose tissue blood flow decreased from rest to after sprint 1 in all 3 subjects, then gradually increased until the end of the experiment, 2 hours after the last sprint (Figure 3). Each data point for flow measurements (pre-exercise, during the rest period between each of the three sprints, and at 2 hours after the last sprint) represents the mean of repeated flow measurements over 15 minutes at steady state.

**Figure 3.**
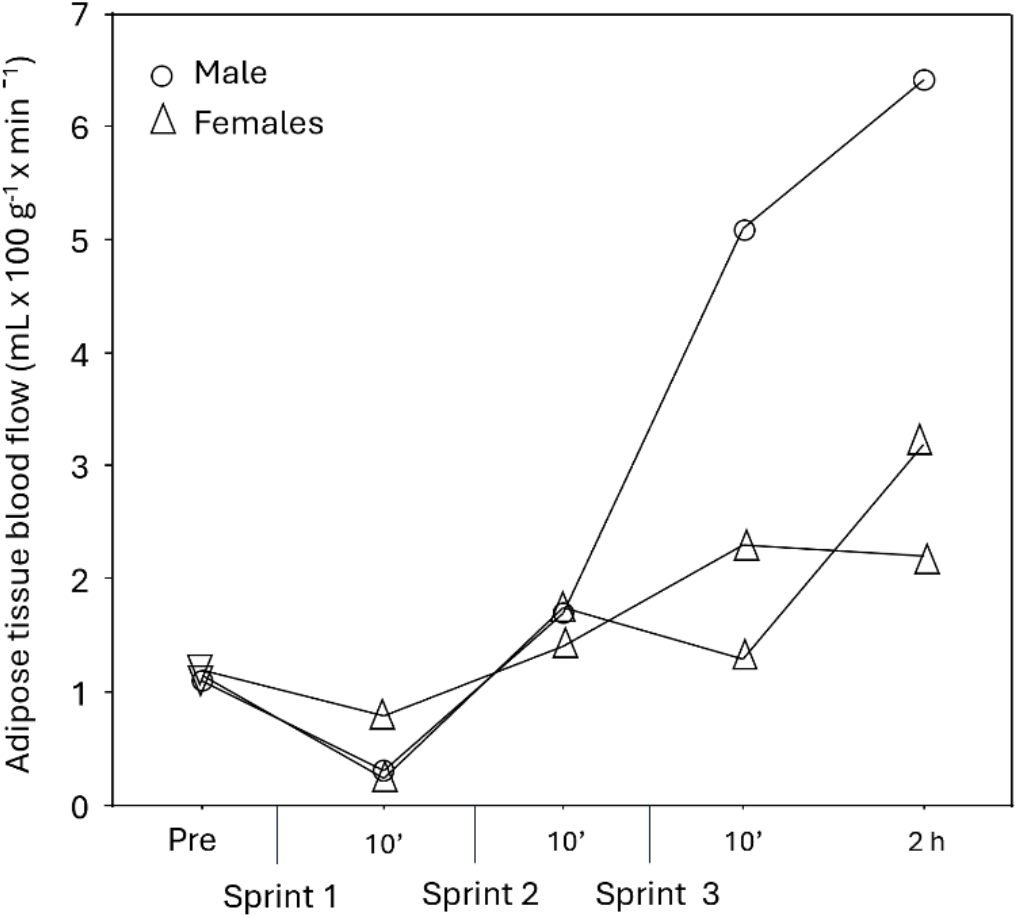
Abdominal subcutaneous adipose tissue blood flow pre-exercise, during the exercise session (3 × 30-s sprint with 20 minutes rest in between) and 2 hours after the last bout of exercise in 3 subjects. Each data point in the figure represents the mean of repeated flow measurements over 15 minutes at steady state.

### Adipose tissue differential gene expression

To test our first hypothesis, that acute SIT induces an increase in post-exercise adipose tissue gene expression related to enhanced lipolysis and energy expenditure, differentially expressed genes (DEGs) were analysed from the microarray. In the SIT group of 12 subjects, 34 differentially expressed genes were identified, with 13 upregulated (e.g. *MIR604, MIR938, ST6GALNAC3 and BMP4*) and 21 downregulated (e.g. *CTGF (CCN2), PER1, NEDD9* and *NFKBIA*), with an adjusted P < 0.1 (Figure 4, Table 2, Table S1). In the CON group of 6 subjects, none of the genes met the cut-off of adjusted P < 0.1, although fold changes for most of the DEGs in the SIT group exercise were of similar direction and magnitude as in the CON group (Figure 5, Table 2, Table S2). Interactions (time x load) were found for only 15 genes, of which 8 were uncharacterised (Table S3). Among the characterised genes, *GHR* was notable, as the secondary hypothesis was that the exercise-induced response of specific GH-responsive genes and the exercise-induced increase in serum GH were associated. Due to the few interactions observed, DEGs were also analysed independently of loading condition in all 18 subjects, where 74 differentially expressed genes were found, with 30 upregulated and 44 downregulated at an adjusted P < 0.05 (Table 3, Table S4). Some of the most downregulated genes were clock genes or genes regulated by clock genes, such as *PER1, NR1D1* and *CIART*, and the clock-regulated gene CTGF (Figure 6, Table 3), all of which decreased independent of SIT or CON (3-factor ANOVAs: time, load and sex; unadjusted P < 0.001 for time for all four genes). Adjusted P-values for the downregulation of *PER1, NR1D1, CIART*, and CTGF independent of group from *limma* were P < 0.001, P = 0.092, P = 0.005 and P < 0.001, respectively (Table 3, Table S4).

**Table 2.**
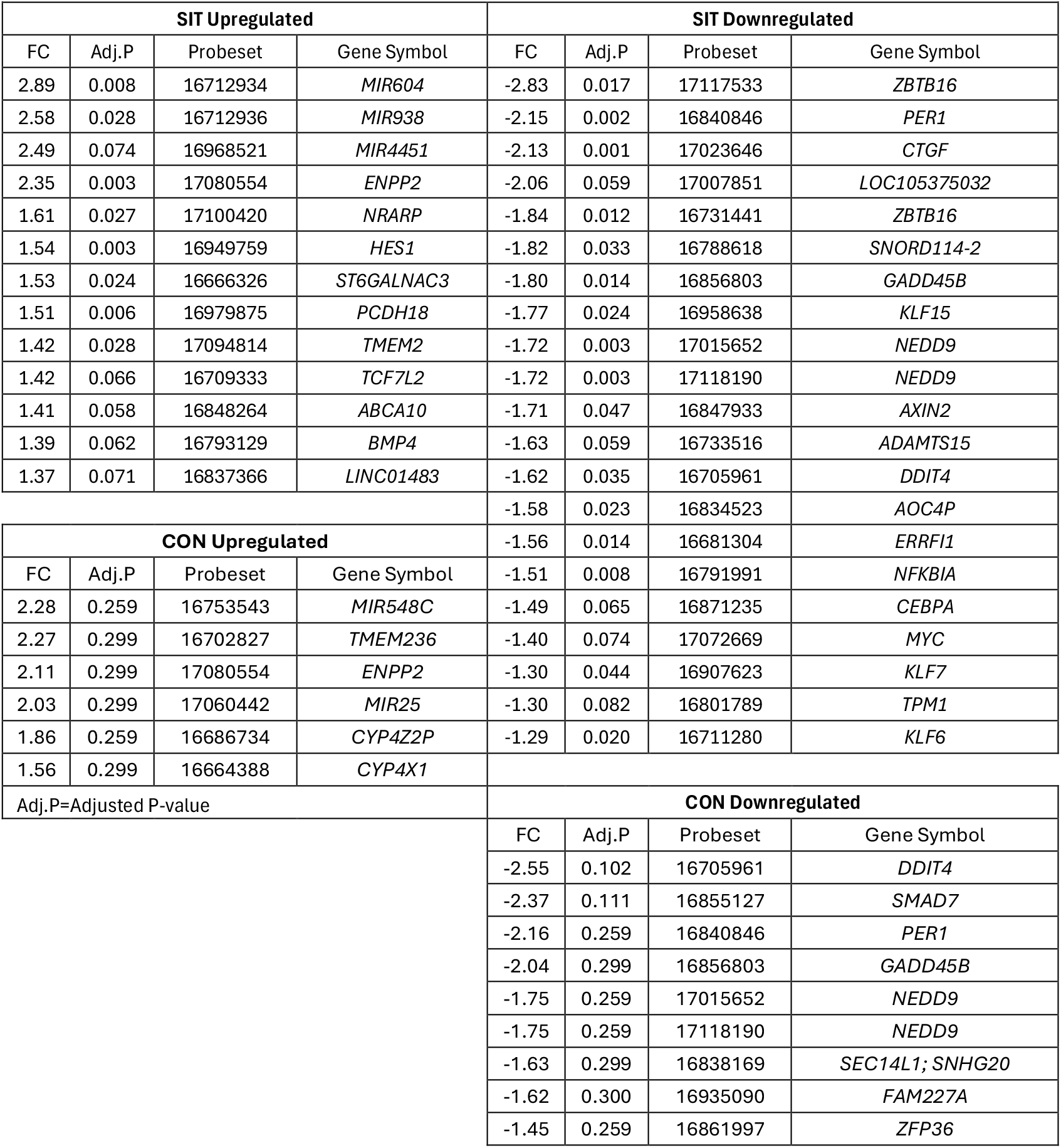
Differentially expressed genes expressed as fold change (FC; post/pre; FC < 1 is expressed as neg FC) in subcutaneous adipose tissue in SIT (6 females and 6 males) and CON groups (3 females and 3 males). Data was analysed by *limma* and adjusted P < 0.1 for SIT and P < 0.3 for CON. © 2000-2025 QIAGEN. All rights reserved.

**Table 3.**
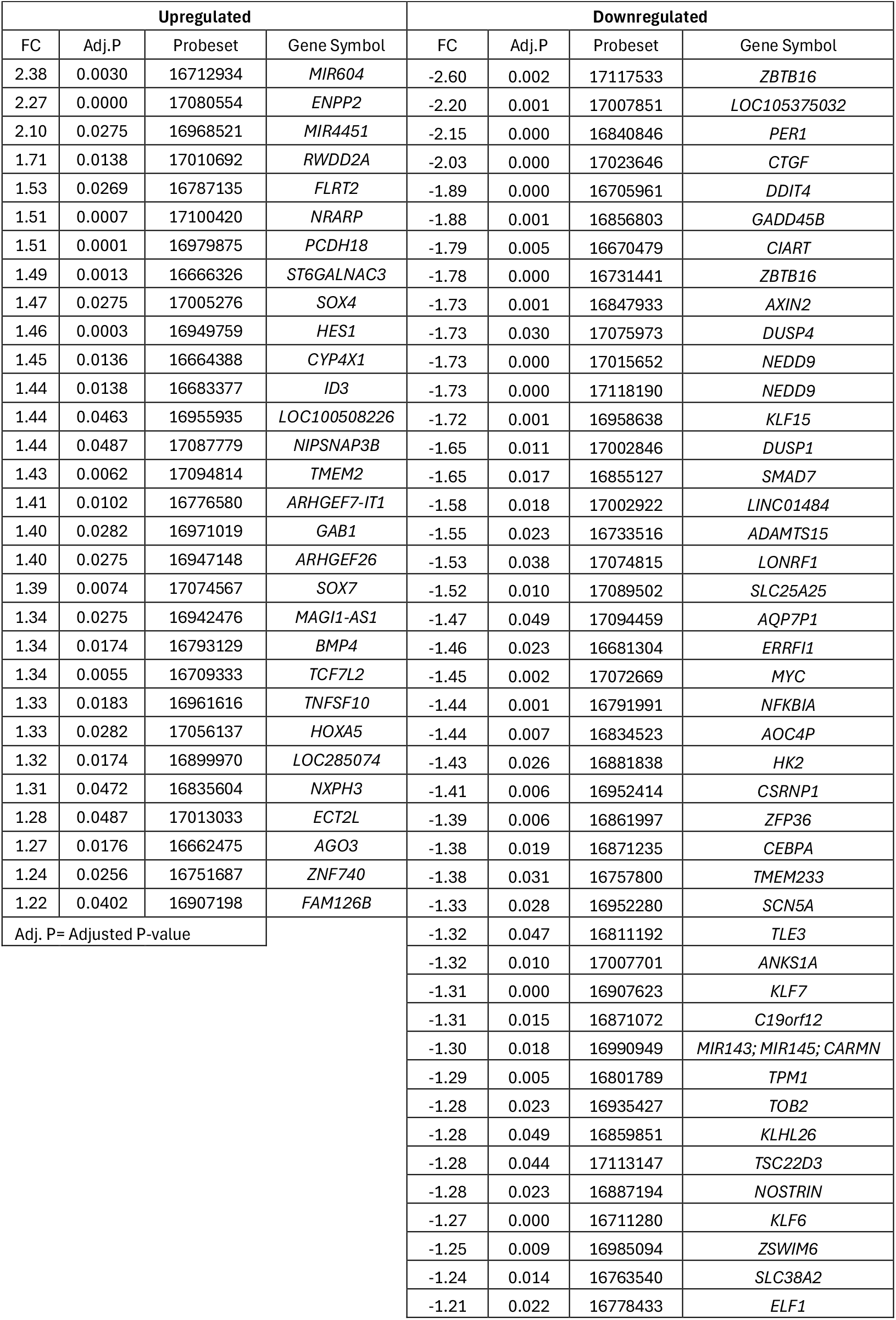
Differentially expressed genes expressed as fold change (FC; post/pre, FC < 1 is expressed as neg FC) in subcutaneous adipose tissue in 9 females and 9 males. SIT and CON combined. Data was analysed by *limma* and adjusted P < 0.05. © 2000-2025 QIAGEN. All rights reserved.

**Figure 4.**
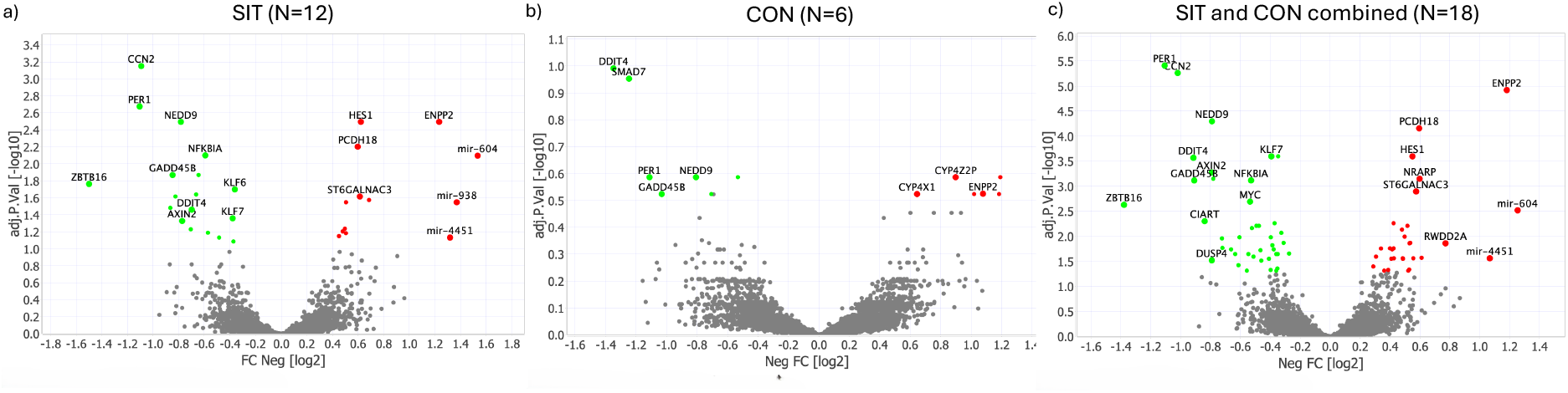
Volcano-plots a) SIT (12 subjects; cut off at adjusted P < 0.1), b) CON (6 subjects; cut off at adjusted P < 0.3), c) SIT and CON combined (18 subjects; cut off at adjusted P < 0.05) showing log2-fold change (FC Neg) for post-exercise/pre-exercise (x-axis) along with -log10 adjusted P-value (y-axis) from the differential gene expression analysis with *limma*. © 2000-2025 QIAGEN. All rights reserved.

**Figure 5.**
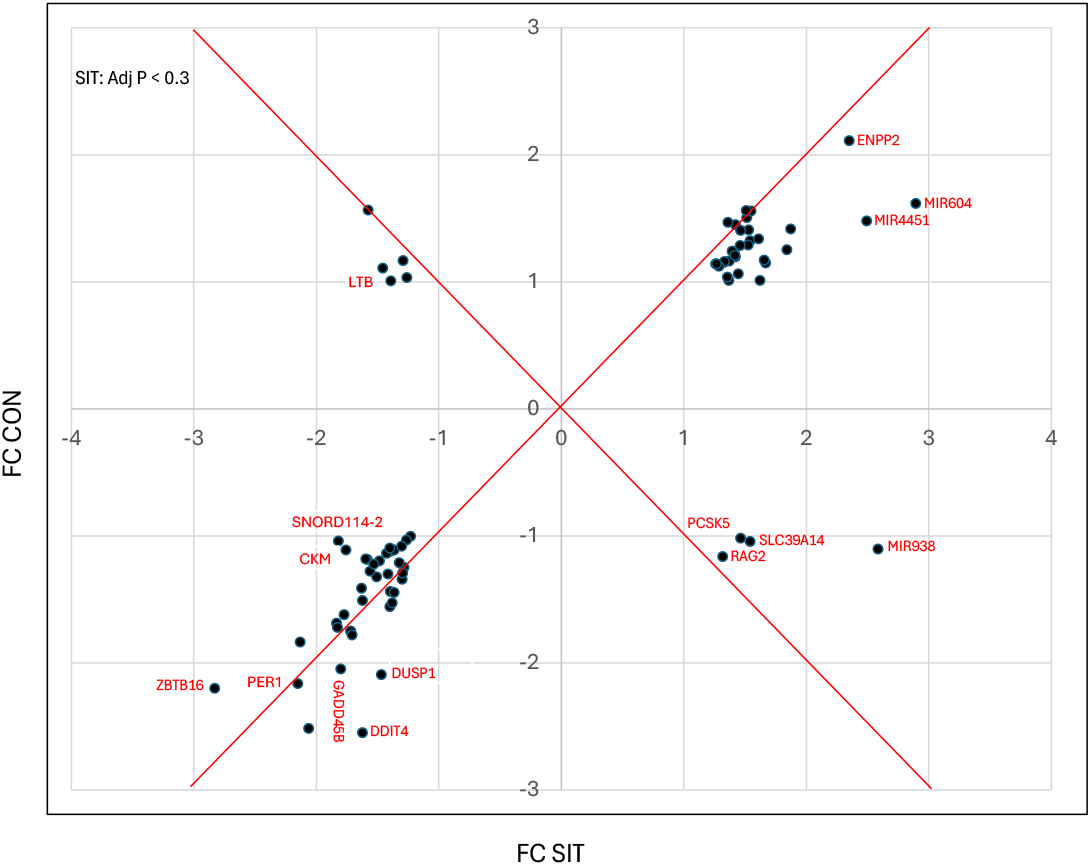
Exercise-induced fold changes (FC) in SIT (12 subjects) versus CON (6 subjects). The 77 included genes were obtained at a cut-off of adjusted P < 0.3 for the SIT group in the *limma* analysis.

**Figure 6.**
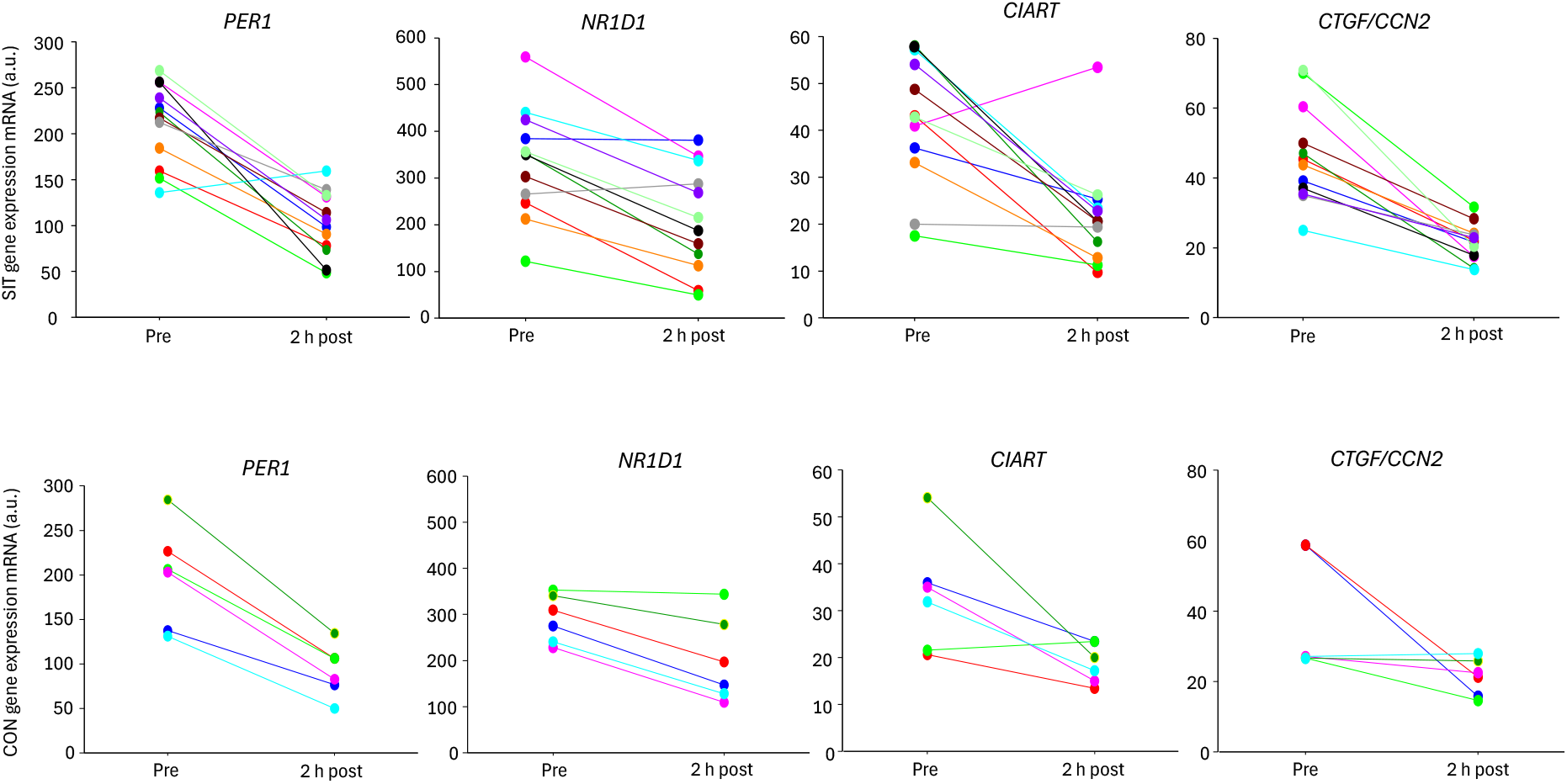
Individual values of core clock gene expression *PER1, NR1D1* and *CIART* and of *CTGF*, a downstream target of core clock genes, pre-exercise and 2 hours after the last bout of exercise in SIT (12 subjects) and CON (6 subjects).

### Predicted anti-inflammatory response

Predicted change (2 hours post-exercise – pre) in activity state was identified by IPA Upstream Analysis. In the SIT group, a list of 77 characterised genes with adjusted P-value <0.3 was analysed by IPA with a z-value cut-off at ±2 and P-value cut-off at P < 0.05 (Table S5). Only a few factors were predicted to be activated, for example, the transcription regulators SOX2, PAX5 and TCF12, and the kinase TGFBR2. However, several factors were predicted to be inactivated, such as the transcription regulators CREB1, LDB1, KLF6, SMAD3, NFKB1 and RELA, as well as cytokines including CSF1, CSF2 and TNF. Further examples of inactivated factors are the growth factors EGF and PDGF-BB, the kinases PI3K and p38MAPK, the clock protein BMAL1 (z = 1.95), and the hormones aldosterone, progesterone and insulin.

In the CON group, a list of 136 characterised genes with ±1.6 FC was also analysed by IPA with a z-value cut-off at ±2 and P-value cut-off at P = 0.05 (Table S6). As in the SIT group, only a few factors were upregulated. Among the inactivated factors, several were found to be regulated in both SIT and CON, such as SMAD3, NFKB1, CSF1, EGF, p38MAPK, aldosterone and insulin. This finding is consistent with only a few interactions in DEGs between SIT and CON. The pattern for the predicted changes of various factors in AT is consistent with decreased inflammation (e.g. NFKB1 and cytokines) and decreased adipogenesis (TGFBR2). Upstream Analysis (z-value cut-off at ±2 and P-value cut-off at P = 0.05) was also performed on the pooled data from a list of 110 characterised genes, P < 0.1 (Table S7) The outcome was similar to the separate group analysis, with few activated factors (e.g. TGFBR2 z = 2.4, P = 7 × 10^−3^and GH z = 1.8, P = 2 × 10^−10^) and several inactivated factors indicating decreased inflammation, such as NFKB1, TGFB1 and various cytokines, growth factors and hormones.

### Associations between adipose tissue gene expression and growth hormone

To test our secondary hypothesis regarding associations between exercise-induced gene expression and GH, we performed regression analyses. First, we conducted a global regression analysis between post-exercise gene expression (adjusted for pre-exercise values) and serum GH peak (adjusted for pre-exercise values). Both SIT (all-out bouts) and CON (unloaded exercise) were included (n = 18). The genes with the lowest adjusted P-values were *ARHGAP24* (P = 0.073), *TAF4B* (P = 0.114), *ARID5B* (P = 0.114), *MIR604* (P = 0.114), and *MIR938* (P = 0.114), all of which were directly related to the GH peak (Table S8). Second, we performed a multiple regression analysis on the pre-selected GH-targets *CISH, PTEN* and *G0S2* and on the genes identified from the global regression above. The exercise-induced changes in *ARHGAP24, ARID5B, TAF4B, MIR604, MIR938, CISH*, and *PTEN* mRNA were found to be directly and *G0S2* mRNA inversely related to the exercise-induced increase in GH (GH peak – pre) (Figure 7). The interaction term [GH peak – pre] x load increased the explanation value (R^2^) significantly for the exercise-induced change in gene expression for *ARID5B* (P = 0.052), *MIR938* (P = 0.016), *CISH* (P = 0.036), and *PTEN* (P = 0.030), and therefore separate regressions for SIT and CON were presented for these genes (Figure 7). For a given exercise-induced increase in GH, the exercise-induced change in gene expression was higher in SIT than in CON (Figure 7). For *ARHGAP24, MIR604, TAF4B*, and *G0S2* mRNA, individuals in the SIT and CON groups followed the same regression line, but individuals in the CON group were found in the lower range of the X-variable GH peak – pre (Figure 7). Overall, this indicates a greater response in the SIT group than in the CON group for all the GH-related genes described above.

**Figure 7.**
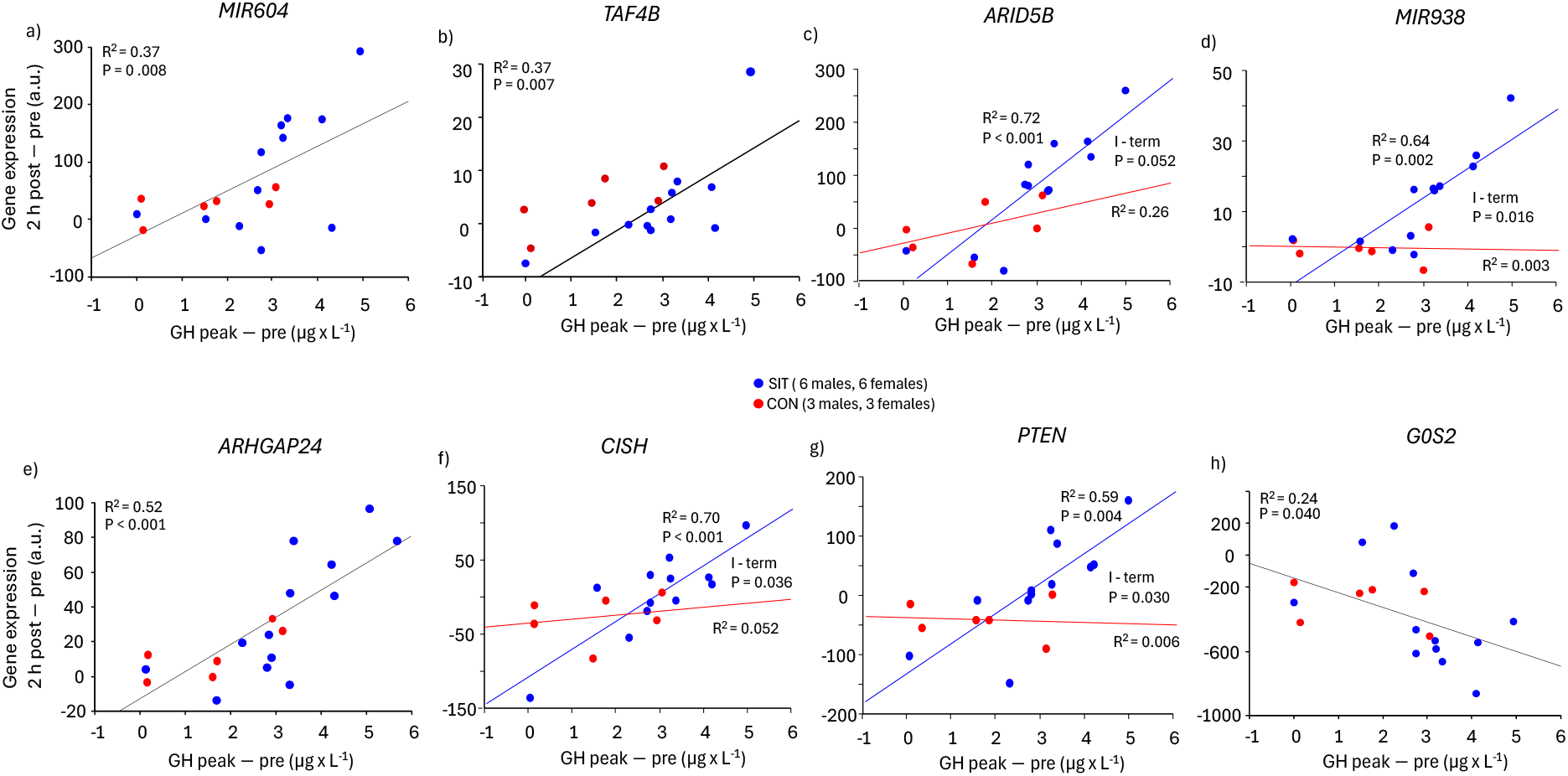
Exercise-induced changes of gene expression (2 hours post – pre) related to increase in serum GH (GH peak – pre) in SIT (12 subjects) and CON (6 subjects) groups combined. The genes presented were those identified from the genome-wide regression analysis (a-e) and the pre-selected GH-targets (f-h). Multiple regression analyses (Y = gene expression 2 hours post – pre, X1 = GH peak – pre, X2 = load (SIT or CON), X3 = interaction (GH peak – pre) x load were performed to see if SIT and CON subjects followed separate or the same regression line. In case of a significant interaction (GH peak – pre) x load between SIT and CON, the interaction P-value and separate R^2^ for SIT and CON are presented. In case of no interaction, the single regression R^2^ is given. Serum GH (GH peak – pre) data were log2-transformed.

To analyse whether load or sex contributed to interindividual variation in the seven GH-associated genes described above, 3-factor ANOVAs (time, group, sex) were performed. Time x load interactions were found for *MIR938* (P = 0.006), *CISH* (P = 0.046), *TAF4B* (P = 0.060), and *PTEN* (P = 0.084), or time x load x sex for *MIR604* (P = 0.114) (Figure S2).Based on the outcome of the 3-factor ANOVAs, 2-factor ANOVAs were analysed separately for the SIT group. Sex x time interactions were found for *MIR604* (P = 0.030), *MIR938* (P = 0.038), *CISH* (P = 0.003), and *PTEN* (P = 0.140), with males showing increased expression for *MIR604* (P = 0.008), *MIR938* (P = 0.003), *CISH* (P = 0.004), and *PTEN* (P = 0.094), but not females (unadjusted P-values). These interactions indicate that both loading conditions and sex contributed to the interindividual variations in these GH-associated genes.

## DISCUSSION

### Key findings

Circulatory growth hormone (GH), which increases lipolysis in abdominal subcutaneous adipose tissue (AT) (34) demonstrated a greater response to acute sprint interval exercise (SIT) than to unloaded bouts of exercise (CON). Despite this, only a few interactions were found in the differentially expressed genes (DEGs) in AT between SIT and CON in the genome-wide approach. However, one of these genes was the growth hormone receptor (*GHR*), indicating a GH-effect on AT that differed between SIT and CON. Nevertheless, we found no major post-exercise enrichment of DEGs related to lipolysis or thermogenesis in AT. Yet, the exercise-induced changes in gene-expression of the pre-selected GH-targets *CISH, PTEN* and *G0S2* were associated with the exercise-induced increase in serum GH. This supports a GH-mediated effect of SIT on lipolysis and thermogenesis. In the genome-wide approach, the exercise-induced changes in gene-expression of *ARHGAP24, TAF4B, ARID5B, MIR604* and *MIR938* were associated with the increase in serum GH. The function of these genes is not well understood, but some relationship to AT metabolism has been demonstrated for ARID5B (35) and MIR604 (36, 37). Moreover, the upstream analysis of the DEGs suggested an anti-inflammatory response to acute SIT. Surprisingly, a similar trend was also found in CON. Among the most downregulated genes, were the core clock genes *PER1, NR1D1* and *CIART*, which decreased similarly in both SIT and CON. Finally, some sex-related differences were identified, especially at rest, for serum GH and the GH-responsive gene *CISH* which is consistent with previous studies (16, 18).

### Core clock genes were altered independently of SIT or CON

An important and unexpected finding in this study was that similar fold changes in gene expression over time were observed in both SIT and CON. In particular, we wish to highlight the 2-fold decrease in the core clock gene *PER1* mRNA, which was independent of group. A decrease was also observed for the other two clock genes, *NR1D1* and *CIART* mRNA, independent of group. *PER1* mRNA peaks at 8.30 am and its expression decreases throughout the day (28, 38). Thus, our experiments were conducted during a period when *PER1* mRNA levels decline, which may explain our findings of decreased *PER1* gene expression, independent of SIT or CON. The similar changes in gene expression in SIT and CON in the present study raise important methodological questions. It is possible that the time of day is a stronger stimulus for changes in AT gene expression than SIT itself for certain genes. Indeed, it has been reported that diurnal rhythm is a significant driver of gene expression variations in human AT, with 25% of genes changing their expression over the day due to the diurnal rhythm (28). Clock genes in AT regulate gene expression in energy metabolism, making it essential to control for this in future exercise studies of AT, especially those related to lipolysis and thermogenesis. In fact, in a recent study it was reported that the expression in core clock genes such as *PER1, NR1D1*, and *CIART*, decreased independent of exercise intensity (39). The authors suggested that the non-exercise factors such as fasting or diurnal variations could have influenced upon the exercise-induced response as verified very recently in a BioRxiv preprint (40).

### Role of growth hormone in lipolysis and regulation of gene expression

The exercise-induced increase in growth hormone (GH) has previously been shown to have a delayed stimulatory effect on post-exercise lipolysis (12) and, consequently, most likely also on post-exercise free fatty acids (FFA) turnover, i.e. thermogenesis (9, 10, 41). Our secondary hypothesis was, therefore, that exercise-induced changes in the expression of GH-responsive genes at 2 hours post-exercise were related to the increase in serum GH during the exercise session. The rationale for this was that GH activates the transcription factor STAT5 in AT and subsequently the expression of typical GH-responsive genes (10, 17, 18) in a GHR-dependent manner (16). In the targeted analysis, we selected the genes *CISH, PTEN* and *G0S2* based on their roles in GH-regulated AT lipolysis. CISH is known to be a negative regulator of GH-induced lipolysis, suppressing STAT5 activation to avoid excessive lipolysis (18, 42). GH also regulates lipolysis indirectly by activating PTEN, which in its turn suppresses antilipolytic insulin/Akt signalling (43), and by inhibiting G0S2, which inhibits the enzyme adipose tissue lipase (44). A second reason for choosing these genes was that GH regulates their gene expression in AT, as based on earlier findings from bolus injection of GH into AT (16, 18). We found that the exercise-induced increases in *PTEN* and *CISH* mRNA were directly related to the increase in serum GH, whereas the decrease in *G0S2* mRNA was inversely related to the increase in GH (Figure 7). This is consistent with a GH-mediated effect on post-exercise lipolysis and thermogenesis (9-12).

In the genome-wide analysis between exercise-induced changes in gene expression and the exercise-induced increase in serum GH, the strongest correlations were found for *ARHGAP24, TAF4B, ARID5B, MIR604* and *MIR938* mRNA, all directly related to the increase in serum GH (Figure 7). Interestingly, three of these genes – *ARID5B, MIR604* and *MIR938* – have been found to be upregulated in AT after a bolus injection of GH (16), supporting a causal relationship between the exercise-induced increase in serum GH and changes in their expression. The functions of these genes are not well understood, but some direct relationship to AT metabolism or an indirect relationship through TGFB, which in turn interacts with inflammation and lipolysis, has been suggested (45). ARHGAP25 may reflect decreased TGFB signalling (46), TAF4B interacts with TGFB in osteogenic cells (47), and MIR938 regulates TGFB in pituitary adenomas (48). MIR604 regulates FASN in mouse heart (37). ARID5B is known from mouse experiments to be involved in AT metabolism, with increased expression described as depressing lipolysis and thermogenesis (35). This is opposite of the expected effect, given the direct association between exercise-induced increases in GH and *ARID5B* mRNA. However, AT metabolism is complex, and counter-regulatory mechanisms are necessary to restrict excessive lipolysis (49). This is comparable to the role of CISH in controlling GH-induced lipolysis (42).

### Adipose tissue stress and anti-inflammatory response: Early vs late post-exercise period

We previously reported that genes related to mitochondrial function and fat metabolism in AT were downregulated, while some inflammation markers were upregulated 15 minutes after acute SIT (15). In contrast, at 2 hours post-SIT in the present study, mitochondrial-related gene expression was unchanged, and an anti-inflammatory response was predicted by upstream regulators, with activation of TGFBR2 and inactivation of NFKB1 (36).

Several mechanisms may explain the switch from a potentially dysfunctional AT gene expression pattern in the early post-exercise phase to a more functional pattern later on. At the onset of SIT, there is a rapid increase in sympatho-adrenergic activity (50) and in AT lipolysis (15), while AT blood flow decreases (Figure 3), likely entrapping FFA in AT (supported by falling plasma FFA during exercise and the subsequent post-SIT rebound; Figure 2). This combination of FFA accumulation and reduced AT blood flow likely creates a transient hypoxic, pro-stress environment (51), consistent with early-phase findings of decreased mitochondrial gene expression, hypoxanthine accumulation, and elevated IL-6 in an AT vein (8, 15). Yet, short bursts of AT stress may induce an anti-inflammatory response in AT (52). Possible mediators are IL-6, succinate/SUCNR1 and glutamine as based on findings of the current and our previous studies (8, 15, 53). Both IL-6 (54, 55) and succinate/SUCNR1 (56-58) may promote M1-to-M2 macrophage switching. In the present study, SUCNR1 mRNA increased after SIT (unadjusted P = 0.003; Figure S1) but not after CON and was positively associated with plasma FFA at 2 hours post-exercise (Figure S1). Also, glutamine may act as an anti-inflammatory agent in AT (59). Previously, we reported an increased glutamine to glutamate ratio in AT after acute SIT, most likely driven by ammonia accumulation in AT derived from skeletal muscle during acute SIT (53).

However, an anti-inflammatory AT response was also indicated in CON (Table S2) comparing pre- and 2 hours post-exercise. This suggests that, for example, diurnal rhythms also drive pathways common to those related to exercise (28, 60), and a complex interaction between diurnal rhythms and exercise-specific effects has been demonstrated in mouse AT (60). Nevertheless, the increased IL-6 release from AT and HPX accumulation in AT were related to plasma lactate accumulation and ATP depletion in skeletal muscle as an early response to SIT (8, 15). This implies exercise-specific effects in AT, in addition to other drivers of AT metabolism. An exercise-induced crosstalk between immune cells and adipocytes has recently been demonstrated in humans (61). However, a possible SIT-induced anti-inflammatory response in AT warrants further investigation.

### Earlier studies of acute exercise and AT gene expression

To our knowledge, only one previous publication has examined global AT gene expression after SIT (15). However, there are some studies on the regulation of abdominal subcutaneous AT gene expression at exercise intensities ranging from 30% to 90% of VO2max, reporting between 10 and 4000 DEGs (38, 39, 62, 63), with no obvious relationship between exercise intensity and the number of DEGs. The inflammatory response, as assessed by DEGs, gene set enrichment, or upstream analysis, is inconsistent across these studies. Some reported an inflammatory response (39, 62, 63), while one of them found an anti-inflammatory response (38). None of the previous studies reported upregulation of genes related to lipolysis or thermogenesis.

Differences in study populations, such as obese/non-obese, or sedentary/trained individuals, sex, experimental protocols including exercise modality and intensity, fasting or non-fasting state, timing of post-exercise biopsies, number of biopsies, and time of day, could explain the considerable variation in gene expression outcomes in earlier exercise studies on AT (28, 38, 39, 62-64). The number of DEGs in the current study of non-obese subjects was at the lower end of the range reported in earlier studies of acute exercise on AT. After correcting for background drivers of gene expression, such as time of day, by comparing SIT and CON, an even lower number of DEGs were identified. The lack of non-exercise controls may contribute to the heterogeneous responses observed when comparing different studies.

### Sex differences in gene expression at rest and in response to SIT

In the present study, an equal number of males and females were included, although there were no specific sex-related research questions. The statistical power was likely too low to identify sex differences in DEGs, especially given possible interference from diurnal variations in AT gene expression. However, sex is an important determinant of AT metabolism at rest, after a meal, or during exercise (65-67).

Regarding SIT, several sex-related systemic changes could potentially affect the AT transcriptome. For example, females reached the level of GH peak faster than males during SIT in the present and previous studies (7, 27). Moreover, in females, elevated serum GH levels were observed already pre-exercise in both SIT and CON, possibly being an anticipatory response to exercise (7, 20, 24-27, 68). In line with this sex-related difference in serum GH profile, the GH-responsive gene *CISH* (23) also showed several-fold higher expression pre-exercise in females than in males (Figure S2). This suggests that the sex difference in serum GH is also reflected at the gene transcriptional level.

Subjects were randomly assigned to SIT or CON just prior to the onset of exercise, which might explain an anticipatory response in both SIT and CON. The response to exercise also differed between males and females for *CISH* mRNA. In the SIT group, males showed an increase and females a decrease in *CISH* mRNA (time x sex; unadjusted P = 0.003). A similar pattern was observed for the GH-associated genes *MIR604* and *MIR938* (time x sex; unadjusted P = 0.030 and P = 0.038, respectively).

### Acute response to SIT – skeletal muscle vs adipose tissue gene expression

Exercise-induced changes in gene expression in skeletal muscle were not analysed in the present study but were examined in a previous study using a similar SIT protocol as in the current one (27). We found a clearly divergent response of skeletal muscle to acute SIT compared to that in AT, consistent with an earlier report (38). The number of DEGs was more than 10-fold higher in skeletal muscle than in AT. Examples of activated upstream regulators in the skeletal muscle include factors related to inflammation, such as transcription factors, cytokines, growth factors, and insulin. In AT, such factors were predicted to be inactivated (e.g. SMAD3, NFKB1, TNF, EGF, and insulin). One factor in common to both skeletal muscle and AT, was the predicted activation of GH. A discordant pattern between adipose tissue and skeletal muscle gene expression was also observed after strength exercise combined with endurance training (69). The divergent response of skeletal muscle and AT was not unexpected, given the profound differences in the function of these two tissues. However, some associations in the acute response of energy-related metabolites between these tissues have been described previously (15). One possible common factor is the systemic hormonal response. One example is the response of *CISH* mRNA to acute SIT. Females had higher pre-exercise levels than males in both tissues, similar to the pronounced sex difference in serum GH in a pre-exercise situation (7). Regarding the response of *CISH* mRNA to SIT, males had a larger response than females in both skeletal muscle (27) and AT (Figure S2).

### Strength and limitations

A strength of the present study is the inclusion of a control group with unloaded exercise (CON). This is important to distinguish the effect of SIT exercise from effects of other drivers of gene expression, such as diurnal rhythms or fasting in AT. A limitation is that the CON group may have been underpowered. Based on the results, the CON group should have been the same size as the SIT group. We did not expect changes of the same magnitude as in the SIT group, and to minimise the number of subjects undergoing invasive procedures, we considered 6 subjects a reasonable sample size for the CON group. Only two biopsies were performed, which limits the temporal resolution in gene expression; however, it is also a strength to avoid a repeated-biopsy effect that might induce an inflammatory response (62). It is also a strength that we included a balanced number of males and females in the study, although this may increase interindividual variability in gene expression. Furthermore, we chose to study a young, healthy, non-obese population, similar to the population in which SIT-induced post-exercise energy expenditure was demonstrated (5). Finally, a notable strength is the use of a well-characterised experimental protocol for the response to SIT, as applied in our previous studies (15).

## CONCLUSION

Following acute sprint interval exercise (SIT), we found no major enrichment of differentially expressed genes related to lipolysis or thermogenesis in abdominal subcutaneous adipose tissue (AT). However, an interaction was observed between the exercise-induced change in *GHR* mRNA between SIT and controls with unloaded exercise (CON), supporting the demonstrated associations between exercise-induced changes in GH-responsive genes in AT and circulatory GH. This partly confirmed our secondary hypothesis. Nevertheless, the observation of changes in AT gene expression also in CON raised important methodological concerns. Clock genes in AT are known to regulate genes involved in energy metabolism (28). Sex was also found to influence both the circulatory GH response to exercise and, for example, the well-known GH-responsive gene *CISH*. This underscores the necessity to control for both diurnal variations and sex in future exercise studies on the regulation of lipolysis and thermogenesis in AT.

## AUTHOR CONTRIBUTIONS

ME, EJ, BN and JB conceived and designed the research. ME, EJ, BN, HR, TÖ and JB performed experiments. EJ, ME and BN and JB interpreted the results of experiments. BN and TÖ performed biochemical analysis and RNA preparations. ME and EJ analysed data. ME, EJ and HR prepared the figures, and HR submitted raw data to GEO. EJ and ME drafted manuscript. EJ, ME, BN, HR and JB edited and revised the manuscript. All authors approved the final version of manuscript.

## ACKNOWLEDGMENT

We thank all study participants and acknowledge the excellent technical support provided by Marika Rönnholm and Ashwini Gajulapuri (Karolinska Institutet). We also appreciate the professional bioinformatics support of David Brodin and thank the core facility at BEA, Bioinformatics and Expression Analysis, located at NEO, which is supported by the Board of Research at Karolinska Institutet and the Research Committee at Karolinska University Hospital. We are also grateful to Eric Rullman (Karolinska Institutet) for his expert assistance with bioinformatic analysis. Finally, we thank Tommy Lundberg (Karolinska Institutet) for his valuable feedback during manuscript preparation.

## DATA AVAILABILITY STATEMENT

Data are available upon request from the corresponding author.

## ETHICS STATEMENT

Subjects were fully informed about the procedures and potential risks of the experiment before giving written informed consent to participate in the study, which was approved by the Ethics Committee at Karolinska Institutet (Dnr 2015/574-31, 2015/1054-32, 2015/1646-32, 2015/2340-32).

---

**Figure S1.**
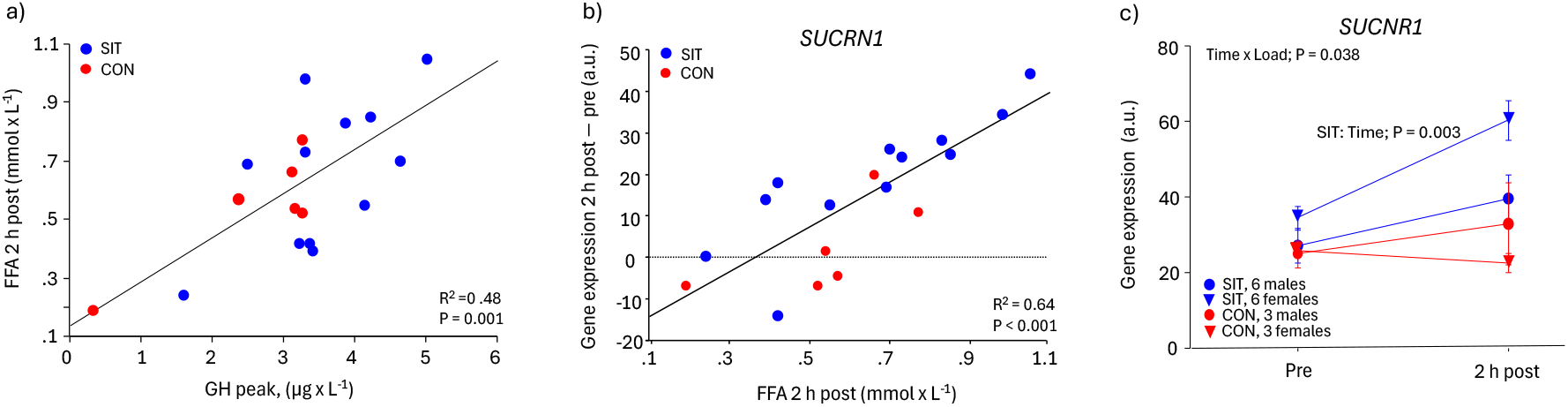
a) Plasma free fatty acids (FFA) 2 hours post-exercise related to serum GH peak, b) exercise-induced changes of SUCNR1 gene expression (2 hours post exercise – pre exercise) related to increase in serum GH (GH peak – pre) in 18 subjects with SIT (all-out bouts) and CON (unloaded exercise) and c) SUCNR1 gene expression pre-exercise and 2 hours post-exercise divided by load (SIT, 6 females and 6 males) and CON, 3 females and 3 males) and sex. SUCNR1 data were log2-transformed and are mean ± SE.

**Figure S2.**
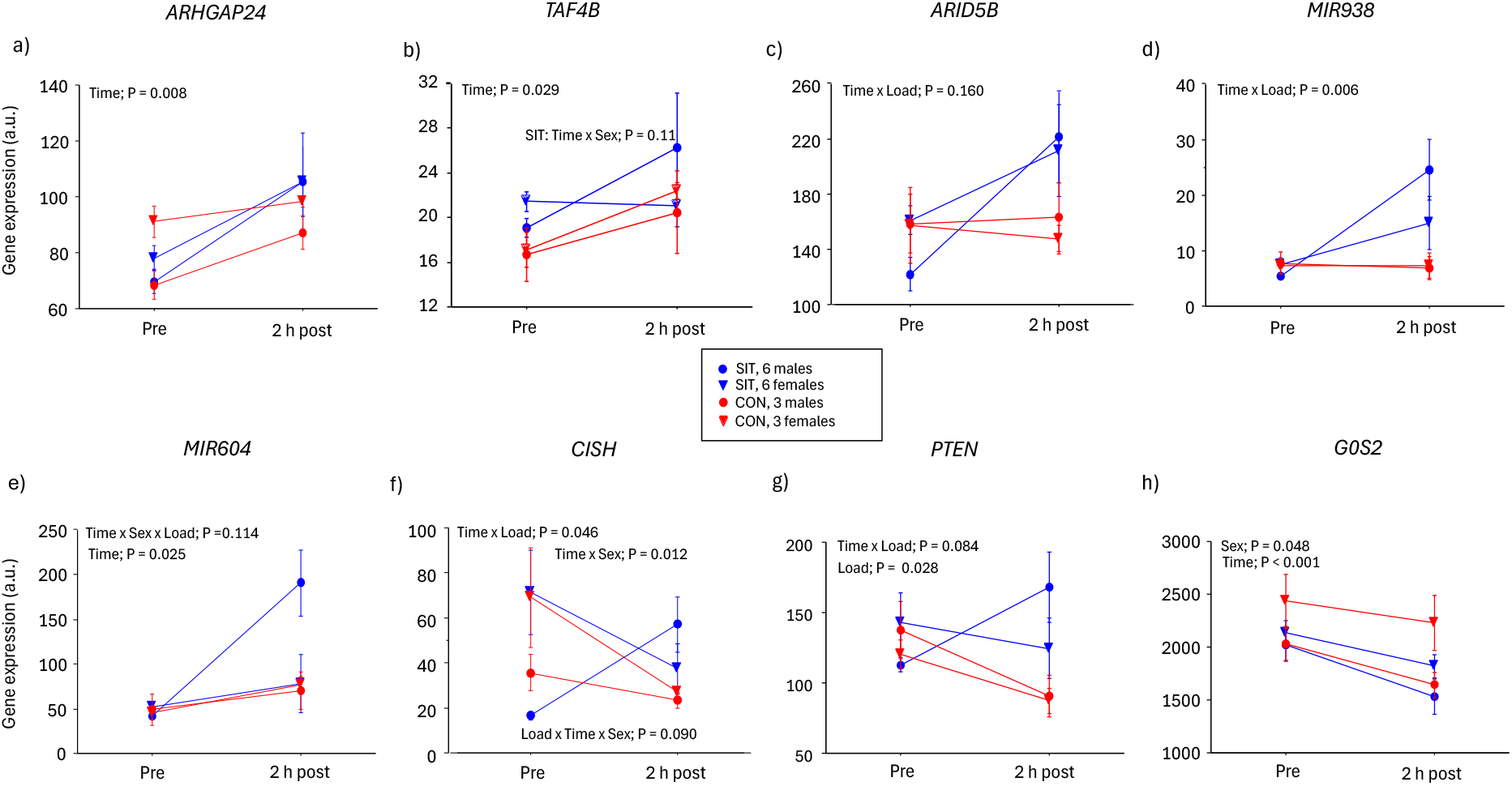
Gene expression pre-exercise and 2 hours post-exercise divided by load (SIT, 6 females and 6 males) and (CON, 3 females and 3 males) and sex. The genes presented were identified from the genome-wide regression analysis (a-e) and from the pre-selected GH-targets (f-h). Serum GH (GH peak – pre) data were log2-transformed. Values are mean ± SE.

